# Lactose intolerance: Magnitude and Associated factors among malnourished under-five Children in Yekatit 12 Hospital Medical College, Addis Ababa, Ethiopia

**DOI:** 10.1101/2020.09.17.301135

**Authors:** Birhane Gebremariam, Abebe Edao, Tigist Bacha, Mistire Wolde

**Affiliations:** Department of Medical Laboratory Science, College of Health Sciences, Aksum University; Department of Medical Laboratory Sciences, College of Health Sciences, Addis Ababa University, Ethiopia; Department of pediatrics emergency and critical care, College of Health Sciences, Addis Ababa University, Ethiopia

**Keywords:** lactose intolerance, Magnitude, lactase deficiency

## Abstract

**Background:** Lactose intolerance (LI) is a pathological condition characterized by the inability to digest sugar, lactose, due to the absence or insufficient activity of the lactase enzyme (β-galactosidase). Currently, laboratory diagnostic procedures for LI are poorly practiced in Ethiopia, and so LI treatment is based on an empirical diagnosis. Thus, the **objective** of the study was to determine the magnitude and associated factors of lactose intolerance among malnourished under-five children in Yekatit 12 Hospital Medical College, Addis Ababa, Ethiopia from March 2018 – July 2018.

**Methods:** A cross-sectional study was conducted among malnourished under-five children admitted in the pediatric unit, Yekatit 12 Hospital Medical College, Ethiopia. By using a convenient sampling technique structured questionnaire was administered to gather information on the socio-demographic characteristics of study participants and associated risk factors of LI. Moreover, a fresh stool sample was collected from the study participants to measure stool pH, reducing substances, and microscopy examination for intestinal parasites. Data entery and analysis was done using the Statistical Package for Social Sciences (SPSS 21) software.

**Results:** The study included 169 malnourished under-five children. Among those 90 (53.3%) were male with a median age of 12 months. The magnitude of LI was 18.3%. Among the study participants, the highest numbers of LI cases were in the age group of less than 12 months; 17(10.1%) followed by 13-24 months; 13(7.7%). Factors that show significant association with LI on bi-variate logistic analysis were family history of lactose intolerance (P=0.043) and diarrhea (P=0.001). In addition; the problem after taking milk (P=0.007), type of therapeutic milk formula (P=0.001), and frequency of stool/24hr (P=0.023) were found to be independent predictors factors of lactose intolerance in the study population.

**Conclusion:** The magnitude of LI was high in the study setting. Thus, more attention should be given on the proper laboratory diagnosis of LI, for better management of cases at the Yekatit 12 hospital. In addition, similar large scale studies at the molecular level are required to strengthen the present findings of LI in Ethiopia.

## Background

Lactose, the major carbohydrate found in milk and dairy products, is a disaccharide composed of two simple sugars, glucose and galactose joined by a glycosidic linkage as β-galactose 1, 4 glucose (1,2,3).Lactose intolerance (LI) was first described by Hippocrates around 400 (B.C) years. However, LI has been recognized and diagnosed as a medically important disease only in the past 50 years (4, 5). LI is a pathological condition characterized by the inability to digest lactose due to the absence or insufficient activity of lactase enzyme (β-galactosidase) (6, 7).

Lactose malabsorption (LM) or hypolactasia is the most common type of carbohydrate malabsorption and is caused by low lactase levels. Lactose intolerance occurs when the malabsorption causes symptoms (i.e. diarrhea, abdominal discomfort, flatulence, and bloating). These symptoms after lactose ingestion are known as lactose intolerance. (2,8,9)

Lactose intolerance is a common medical problem, significantly impacts the lives of affected individuals and has limited treatment option (10) .Worldwide, 75% of population loses the ability to digest lactose (11) But this prevalence of LI is strongly linked to age, ethnicity, use of dairy products in the diet and method used for its diagnosis. In populations with a predominance of dairy foods in the diet, particularly northern European people as few as 2% of the population has primary lactase deficiency is common among adults. In contrast, the prevalence of primary lactase deficiency is 50% to 80% in Hispanic people, 60% to 80% in black and Ashkenazi Jewish people, and almost 100% in Asian and American Indian people (1,9,12,13)., In the United States, the prevalence is 15% among whites, 53% among Mexican Americans and 80% in the Black population. The prevalence of primary lactase deficiency is above 50% in Africa (14)

Gastrointestinal symptoms caused due to LI condition induced by milk and milk products malabsorption, indirectly interfere with calcium dietary intake. Thus, LI has be,en considered as a risk factor for low bone mineral density, osteoporosis, and depression. Symptoms of LI can affect patients physically, psychologically, and socially (15-18)). More than one-third of all study subjects had gastrointestinal symptoms after lactose ingestion (15) Even though lactose may be a beneficial nutrient for undernourished children; research is needed to define the balance between beneficial and detrimental effects of lactose in undernourished children at different ages and with different degrees of diarrhea and intestinal integrity (19, 20).

There are different laboratory diagnostic methods for LI including determination lactase activity of a jejunal biopsy, lactose tolerance test, hydrogen breath test (HBT), analysis of faecal pH and reducing substances and genetic studies (9, 12,21,22)

Currently in Ethiopia LI treatment is based on an empirical diagnosis. Treatment based diagnostic approach is being utilized to diagnose whether children are lactose intolerant or lactose tolerant. This is done by observing for adverse and poor response in the children who are taking therapeutic milk formula. Laboratory diagnostic procedures for LI are poorly practiced in Ethiopia. Furthermore, recently there is no study conducted on the magnitude and associated factors of lactose intolerance among malnourished under-five children in Ethiopia, where LI is more significantly expected to be seen. Thus, it is difficult to discuss exactly LI situation in Ethiopia. Therefore, the aim of this study is to determine the magnitude and associated factors of LI among under-five malnourished under-five children. Thus, we aimed to determine the magnitude and associated factor of lactose intolerance among malnourished under-five children in Yekatit 12 Hospital Medical College, Addis Ababa, Ethiopia.

## Material and Methods

The present study was a cross-sectional study, conducted from March - July 2018 in the pediatric unit of Yekatit 12 Hospital Medical College, Addis Ababa, Ethiopia. Addis Ababa is the capital city of Ethiopia. Addis Ababa lies at an altitude of 7,546 feet (2,300 meters) above sea level, located at 9°1′48″N 38°44′24″E Coordinates: 9°1′48″N 38°44′24″E. Based on the 2007 census conducted central statistical agency of Ethiopia Addis Ababa has a total population of 2,739,551 (23). Yekatit 12 Hospital Medical College is a public hospital found in Addis Ababa city established in 1915. Yekatit 12 Hospital Medical College provides a health care service at an out-patient and in-patient level as a referral hospital center for health centers and hospitals in Addis Ababa as well as different regions of the country. It is one of the centers serving as a nutrition therapy. The Pediatric unit has 8 beds for the admission of malnourished under-five children. The hospital is using the Federal Ministry of Health guideline for the management of severe acute malnutrition adapted from the United Nations Children’s Fund (24).

As a source of population, all children who visit the pediatric unit of Yekatit 12 Hospital Medical College during the study period were used. Among them, malnourished under-five children admitted to pediatric unit of Yekatit 12 Hospital Medical College and who were taking therapeutic milk formula during the study period were selected for the study.

As an eligibility criterion for the study participants; children Age (≤5 years), Children with malnutrition, participant who was on therapeutic milk formulas during the study period were considered as an inclusion criterion; meanwhile as an exclusion criterion: Children with known diabetes and Children on proton pump inhibitor, laxative use were used.

During the study, as a dependent Variable Magnitude of lactose intolerance were used, while as an independent variable; Socio-demographic characteristics (age, gender, preterm birth), feeding practice, type of malnutrition, family history of LI, problem after milk is taken, immunization status and Intestinal parasites considered.

The sample size of the study was determined by using the prevalence (11.2%) of prior study on carbohydrate intolerance after acute gastroenteritis in Polish children (25). The sample size was calculated by using the formula for sample size determination for a single proportion. Accordingly, a convenient sampling technique was used to include a total of 169 study participants for the study.

### Data Collection Procedure

After getting approval and permission from Yekatit 12 Hospital Medical College, a structured questionnaire was used to gather information on the socio-demographic characteristics and associated factors with LI. Trained nurse staff collected the demographic data and instructed the family of the children how to collect the stool. A fresh stool sample was collected 24 hours after they start a standard lactose-based therapeutic milk formula. Then, the stool sample was transported from the pediatric ward to the laboratory. Laboratory tests were done in the fresh stool of the participants with measurements of stool pH and reducing substances. Stool pH was tested using pH papers, while the presence of reducing substance was tested by the use of Benedict’s solution. Stool microscopy was also performed to detect intestinal parasite (G.lamblia and E.histolytica).

### Data Analysis and Interpretation

Data were coded, double entered, and analyzed using the Statistical Package for Social Sciences (SPSS 21) software. Categorical variables were summarized as frequencies and percentages, while continuous variables as a median. In the bivariate analysis, odds ratios, 95% confidence interval (CI), and the chi-square test or Fisher’s exact test were used to measure the strength of association between the factors considered and the dependent variable. Multivariate analysis using logistic regression was used to determine the factors that were independently associated with lactose intolerance. Risk factors with p-values below 0.2 on bi-variate analysis were included in logistic regression analysis to identify independent risk factors. P-value < 0.05 was considered for statistical significance. The results was summarized in texts, tables, and bar graphs.

### Ethical Consideration

Ethical clearance was taken from the research and ethics review committee of the Medical Laboratory Science department and Addis Ababa health bureau. In addition, a letter of permission was taken from Yekatit 12 Hospital Medical College. Prior to data collection, written informed consent was obtained from each parent/caregiver of the study participants after explaining the purpose of the study. Confidentiality of the data was maintained by coding of samples and privacy of the respondents was maintained. The result of the laboratory test was communicated to the treating physician/nurse and they could modify the treatment of the participant if they deemed necessary.

### Operational definition

**Lactose malabsorption**: indicates that a sizable fraction of a dosage of lactose is not absorbed in the small bowel and thus is delivered to the colon (26).

**Lactase deficiency**: is defined as markedly reduced brush-border lactase activity relative to the activity observed in infants (27).

**Lactose intolerance**: is considered if fecal reducing substances are ≥0.5g% and stool pH less than 5.5 (28).

**Malnutrition**: refers to deficiencies, excesses, or imbalance in a person’s intake of energy and/or nutrient (29).

## Results

### Socio-Demographic Characteristics

A total of 169 malnourished under-five children admitted in the pediatric unit of Yekatit 12 Hospital Medical College was enrolled in the present study. Among them, 90(53.3%) were male with a median age of 12 months. Moreover, the highest numbers of cases were occurring in the age group of less than 12-month 91(53.8%) followed by 13-24-month 56(33.1%) age group (Table 1).

### Clinical sign-symptom and stool characteristics

Among the study participants, 70(41.4%) had diarrhea; of which 30(42.9%) had persistent diarrhea. Among the study participants, 51(30.2%) had a watery stool, 14(8.3%) had perianal skin erosion and 15(8.9%) had the frequency of stool /24 hours ≥5 times (Table 2). *E*.*histolytica* was found only in 1 patient while *Giardia lamblia* was founded in 8(4.7%) out of the study participants.

### Feeding pattern and malnutrition type of study participants

Among the study participants 156(92.3%) had SAM, 131(77.5%) were on F-75 therapeutic milk formula and 15(8.9%) out of the study participants had experienced problems with cow milk (Table 3).

### Magnitude of lactose intolerance of study participants

According to the criteria set for definition of LI, in the present study the overall magnitude of lactose intolerance was found to be 18.3% (Table 4).

### Factors Associated with lactose intolerance

#### Association between diarrhea characteristics and lactose intolerance

The present study demonstrated that diarrhea and frequency of stool in 24hr show significant association with lactose intolerance while the duration of diarrhea, perianal skin erosion, *Giardia lamblia*, and vomiting does not show a significant association with lactose intolerance (Table 5).

#### Association between baseline characteristics and lactose intolerance

Sex, age, preterm birth and immunization status does not show significant association with LI but family history of LI show a significant association with the lactose intolerance (Table 6). There is a gradual decrease in the magnitude of LI among the age groups (Fig.1).

**Figure 1:**
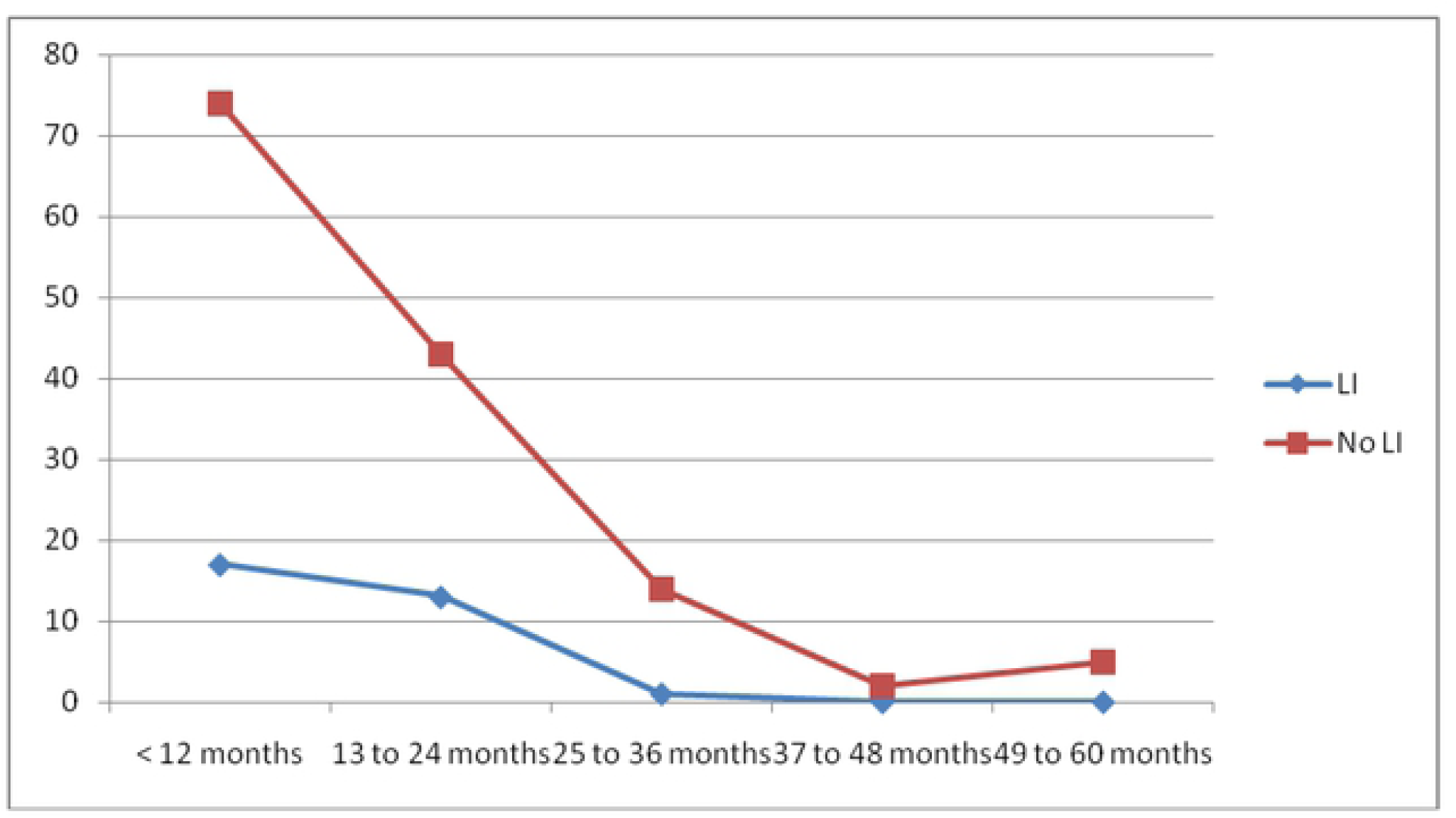
Age distribution by lactose tolerance of malnourished under five children, admitted in the pediatric unit of Yekatit 12 Hospital Medical College from March -July 20I 8.

#### Association between feeding practices and lactose intolerance

In the present study, problem after taking cow milk (P=0.008) and therapeutic milk formula (P=0.004) shows a significant association with lactose intolerance (Table 7).

#### Factors independently predicting lactose intolerance

Factors for the multivariate logistic regression were chosen because they have been shown to be associated with LI and/or they had a P-value of <0.2 on bivariate logistic regression analysis. Accordingly, as shown in Table 8 after multivariate logistic regression analysis; high stool frequency in 24-hour, problem after taking cow milk, and type of therapeutic milk formula were the independent predictors of lactose intolerance. The respective AOR and CI are 3.80 [1.26-11.48, P=0.023], 8.14 [1.78-37.12, P =0.007] and 0.13 [0.04-0.40, P =0.000].

## Discussion

Lactose intolerance is the inability to digest lactose, a disaccharide found in milk, and to a lesser extent in milk-derived dairy products (11). In the present study, the overall magnitude of lactose intolerance was 18.3%. This result wass comparable with a study done in Turkey (16.3%) (16); however is lower than a study done in Uganda (25.5%) (30), India (25.71%) (31), Iraq (41%) (32), South Africa (59%) (33) and Georgia (41.0%) (34). In contrast, it was slightly higher than a study done in Poland (11.2%) (25). This difference might be due to the difference in the diagnostic method used, sample size, study populations. In this study, most of the study populations were malnourished child without diarrhea (58.6%); while most of the other studies included malnourished child with diarrhea. In addition, other difference could be due to difference in study design. This study utilized cross-sectional study design while a prospective cohort study was done in Georgia (34).

In this study 8(25.8%) subjects with lactose intolerance had incomplete immunization status. This finding was nearly similar to a study done in Iraq 9 (21.95%) (32), but lower compared to a study done in Uganda 39(39.8%) (30). In line with this, immunization status had no significant association (P=0.560) with LI which was similar to a study done in Iraq (P= 0.44) and disagree with a study done in Uganda (P< 0.001) (30). This might be explained due to differences in sample size, increased awareness towards immunization utilization, the difference in health policy towards immunization utilization of study area.

In the present study, there was no significant association (P= 0.553) between sex and lactose intolerance. This result was similar to study done in Uganda (P= 0.545) (23), Iraq (P= 0.67) (32). The lack of association with gender might be indicates that if the clinical condition is genetically determined, it is not linked to the sex chromosomes.

In the present study LI was seen in 15 (48.4%) children who had vomiting; a finding lower than a study done in Iraq (75.6%) (32), and in Uganda (70.0%) (30). In this study, there was no significant association (P=0.384) between lactose intolerance and vomiting. This finding contrast with study done in Uganda (P= 0.027) (30), Iraq (P=0.01) (32) and Tbilisi, Georgia (P=0.0011) (34). This might be due to differences in the diagnostic method, study design, and study population. In the present study preterm birth (P= 0.074) had no significant association with LI while the family history of LI (P= 0.043) showed significant association with LI.

Out of the study population, 28.4% had an EBF duration of less than 4 months; which was lower compared to study in Uganda (35.7%) (30), and Iraq which 72% (32). There was no significant association between EBF (P= 0.064) and LI which agrees with a study done in Iraq (P=0.50) (45) but it differs to study done in Uganda (P=0.015) (30).

In the present study LI was seen in 22.6% of the study participants, who had problem after taking cow milk. In addition in the current study, a significant association was demonstrated (P=0.008) between problems after milk taken with LI. This finding was comparable to study done in Tbilisi, Georgia (P= 0.0076) (34); though the result disagrees with a study done in Uganda (50.0%; P=0.175) (30). This could be explained due to differences in the sample size and study population.

In the present study, most of the lactose-intolerant children (58.1%) were on F-75 as compared to 41.9% of children who were in the F-100 therapeutic milk formula. This outcome accords to a study done in Uganda in which most of the lactose-intolerant children (82%) were on F-75 as compared to 18% of children who were in F-100 therapeutic milk formula (30). In line with this, there is a significant association (P= 0.004) between LI and therapeutic milk formula; a finding which contrasts to study done Uganda (P=0.938) (30). This might be due to differences in the study population i.e. there is no equal distribution of study participants among the therapeutic milk formula feeding.

Most children (54.8 %) with lactose intolerance were children less than 12 months of age, which is consistent with a study done in Uganda (68%) (30), Iraq (73.17%) (32), South Africa (69%) (33), and Tbilisi, Georgia (42%) (34). In line to this, there is a gradual decrease in the magnitude of LI from 10.1% in the age group less than 12 months, 7.7% in the 13-24 month age groups, 0.6% in the age group 25 -36 month, and with zero both in the 37-48 and in the 49-60 months age groups; a result consistent to a finding in Uganda (30). This could be attributed to the higher susceptibility of the intestine of children less than 12 months of age; which could be aggravated by the diarrhea and malnutrition. On bivariate analysis age in months had no significant association with LI (P=0.902) a finding similar to a study done in Iraq (P=0.58) (32); but inconsistent with a study done in Uganda (P=0.018) (30)and in South Africa (P=0.047) (33) This could be due to the difference in study subjects in which most of the study participants in the present study were children less than 12 months of age i.e. there was not normal distribution between the age group of study subjects.

Children with LI had higher mean stool frequency (≥4motions in 24 hour period; P= 0.001), a finding consistent with a study in Uganda (30), South Africa (33), and Tbilisi, Georgia (34). This is expected as unabsorbed lactose would remain in the colonic lumen and lead to osmotic diarrhea. Furthermore, undigested lactose may attract fluid, water, and electrolytes into the lumen that the colon cannot handle (5, 6) (1,2)

Seventy (41.4%) of the study participants had diarrhea; of those 42.9% had a diarrheal duration of ≥14days. This seems to strengthen the fact that diarrhea contributes to intestinal mucosal damage associated with variable degrees of malabsorption (1). Among the study population who had diarrhea 31.4% were lactose intolerant (P=0.001) which is lower compared to a study done in South Africa 59% (33), Iraq 41% (32); but higher to a finding in Uganda 25.5% (30). This might be difference in the study population.

In the present study LI was seen more frequently in the children who had persistent diarrhea (43.3%) compared to patients with acute diarrhea (22.5%); this contrasts with a study in South Africa in which duration of diarrhoea was found to be shorter in children with lactose malabsorption than in those without (33). The finding is comparable to study done in Uganda in which LI was seen more frequently in children with persistent diarrhea (34.2%) as compared to (20.0%) with acute diarrhea (30). This confirms the observation that the lactase enzyme is localized to the tips of the intestinal villi, a factor of clinical importance when considering the effect of diarrheal illness on the ability to tolerate lactose. Persistent diarrhoea also results in a more prolonged and extensive damage of the intestinal mucosa and the immature epithelial cells that replace these are often lactase deficient, leading to secondary lactase deficiency and lactose malabsorption (1,35).

At multivariate logistic regression analysis, problem after taking milk (P=0.007), type of therapeutic milk formula (P=0.000) and high stool frequency in 24 hour (P=0.023), were the independent predictors of lactose intolerance in malnourished children. In a study done in Uganda (30), high mean stool frequency (P=< 0.001) was among the factors which independently predict lactose intolerance while in a study conducted in South Africa age of <12 months (P=0.046) was the only factor found to significantly predict lactose malabsorption.

### Strength of the study

This research provided updated information on the current magnitude of lactose intolerance in the study setting which was not done before.

### Limitation of the study

In the present study hydrogen breathe test which is gold standard test was not done because of its unavailability in our country and expensive cost. The Benedict’s test relies on a color change which leads the interpretation of the color to somewhat subjective but this observer bias was minimized by only one researcher conducting the laboratory test. The study was conducted in one hospital and the sample size was small. Besides the study was focused on specific group of patients, malnourished children This could have resulted in decreased variability in the study results. Associated factors with lactose intolerance could be evident if the study was done in larger study sample size. Additionally, culture for bacterial infection and stool antigenic test for Rota virus was not done.

## Conclusion

The overall magnitude of lactose intolerance which is 18.3% among malnourished under-five children in this study setting is high. On bivariate analysis; diarrhea, frequency of stool/24hr, family history of LI, problem with cow milk and therapeutic milk formula show a significant association with LI. In the present study LI was found to be common among malnourished under-five children with diarrhea duration of ≥14days i.e. persistent diarrhea. According to the multivariate logistic regression analysis; high stool frequency in 24-hour, problem after taking milk and type of therapeutic milk formula were found to be the independent predictor factors of lactose intolerance.

## Recommendation

- Lactose intolerance should be considered and routine screening by stool pH and reducing substance would be good if it is done in these hospitals or other health facilities which provide treatment for malnourished children.
- A research with different study design, larger sample size and different variable deserves in this field.
- Malnourished under-five children with higher frequency of stool in 24hr and having problem after taking milk, LI should be considered and managed accordingly
- This study highlights the need for a future study on investigating the impact of the use of the lactose free feeds on the recovery of children with malnutrition whose stool is positive for reducing substances.

